# A versatile microfluidic device for highly inclined thin illumination microscopy in the moss *Physcomitrella patens*

**DOI:** 10.1101/660480

**Authors:** Kozgunova Elena, Gohta Goshima

## Abstract

High-resolution microscopy is a valuable tool to study cellular processes, such as signalling, membrane trafficking, or cytoskeleton remodelling. Several techniques of inclined illumination microscopy allow imaging at near single molecular level; however, the application of these methods to plant cells is limited, due to thick cell walls and necessity to excise a part of the tissue for sample preparation. In this study, we developed simple, easy-to-use microfluidic device for highly inclined and laminated optical sheet (HILO) microscopy using a model plant *Physcomitrella patens*. We demonstrated that microfluidic device can be used to culture living cells and enables high-resolution HILO imaging of microtubules without perturbing their dynamics. In addition, our microdevice enables the supply and robust washout of compounds during HILO microscopy imaging, for example to perform microtubule regrowth assay. Furthermore, we tested long-term (48 h) HILO imaging using a microdevice and visualised the developmental changes in the microtubule dynamics during tissue regeneration. The microfluidic device designed in this study provides a novel tool to conduct long-term HILO microscopy and washout assays using plant cells.

## Introduction

Observing protein dynamics in living cells at a single-molecule level provides researchers with valuable data on protein interactions and their function. Although the popularity of live-imaging approaches continues to increase, several techniques are difficult for plant application, owing to their slow development and challenging optical properties, such as chloroplast auto-fluorescence and thick cell wall^1,2^. One of the key technologies to monitor the cellular events close to the cell surface is total internal reflection fluorescent (TIRF) microscopy. TIRF method is based on the total reflection of illumination beam at the boundary of two different reflective indexes; for example, between coverslip and imaging medium^3^. Therefore, TIRF can be applied only when the molecules of interest are very close to the cell surface, which inevitably limits the TIRF use in plant cells surrounded by thick cell walls. Another similar approach is variable angle evanescent microscopy (VAEM), also known as ‘pseudo-TIRF’, oblique illumination fluorescent microscopy or highly inclined and laminated optical sheet (HILO) microscopy, which allows to image events more distant from the cell boundary^4^. TIRF and HILO share certain similarities and the major difference between them is the angle of excitation beam^5^. For both imaging methods, the sample surface has to be flattened against the coverslip. In animal studies, adherent cells provide a natural system for TIRF imaging; alternatively, several coating agents such as fibronectin or poly-l-lysine can be used to attach cells^3^. However, plant samples are usually prepared by covering the excised tissue with an agar block or second coverslip to increase the visible surface^6^. Tissue excision, as well as limited gas exchange, could cause changes in cellular dynamics over time; hence, the plant samples should be observed soon after preparation. Nevertheless, certain molecules or pathways are active only during particular developmental stages; for example, when the cytoskeleton is dramatically rearranged during the cell division^7,8^. Development of the long-term HILO imaging for plant cells will allow to observe such changes with high resolution.

Microfluidic devices are commonly fabricated from polydimethylsiloxane (PDMS), a biocompatible material suitable for long-term culture conditions^9–12^. PDMS can be moulded into different designs and patterns; for example, a recent study using microdevice has demonstrated that tip-growing plant cells are able to penetrate extremely narrow 1-µm gaps^13^. In addition, a microfluidic nature of the system enables rapid liquid exchange inside the device during live-cell imaging. Microdevice technology has been routinely applied to supply/washout compounds, such as small-molecule inhibitors for *in vitro* or animal cells research.

Here, we test and optimise the compatibility of microfluidic device for HILO imaging in the moss *Physcomitrella patens*, focusing on microtubule dynamics. We demonstrate that our simple, easy-to-use microdevice can be applied in various experiments, such as regular HILO imaging or imaging combined with acute supply/washout of drug compounds. Moreover, we were able to perform a long-term (48 h) HILO imaging using microdevice to visualise the developmental changes in the cortical microtubules array during tissue regeneration.

## Results

### 1. Severe space restrictions cause subtle changes in microtubule dynamics in protonema cells

We designed a simple microfluidic device that includes four independent channels (Figure 1A). This device can be mounted either on a coverslip, for short-term imaging, or on a glass-bottom dish, for long-term imaging. In this study, we used a popular model plant *Physcomitrella patens*^14^, known for its regeneration ability^15,16^. Monthly cultured *P. patens* colonies mostly consist of three tissue types: the protonema comprising tip-growing stem cells, leafy shoots known as gametophores and rhizoids. Diameter of the protonema cells in our culture system is 21.5 ± 2.39 µm (*n* = 16); thus, we hypothesised that the cells grown in shallow channels would be naturally flattened against the coverslip, providing a setting for HILO imaging. We designed and tested three versions of the device with different channel depth: 4.5, 8.5 and 15 µm to determine the optimal conditions (Figure 1A). Initially, we injected small clusters of protonema cells, expressing microtubule marker GFP-α-tubulin^17^, into the device and allowed them to regenerate for 3–4 days (Figure 1B). Notably, we did not observe changes in the viability or growth defects, even in the shallowest 4.5 µm device. Next, we performed HILO imaging of microtubules in regenerated cells. As a reference, we imaged a freshly prepared sample of protonema cells flattened between two coverslips (Figure 1C; Supplemental Movie 1). Although a previous report demonstrated that protonema cells are able to grow in extremely narrow channels^13^, it remained unclear if the space restrictions affect the cellular processes. We quantified microtubule growth velocity (Figure 1D), and found that protonema cells in 4.5 µm device, but not in 8.5 or 15 µm devices, show a slight decrease (0.108 ± 0.026; 0.1324 ± 0.018 and 0.134 ± 0.021 µm/s, respectively [*n* = 10, 9, 8]). Interestingly, we also did not notice any advantages in using 4.5 and 8.5 µm devices in terms of imaging quality or visible cell surface area (Figure 1C; Supplemental Movie 1). Thus, we chose the device with 15 µm channel depth to conduct further experiments.

**Figure 1.**
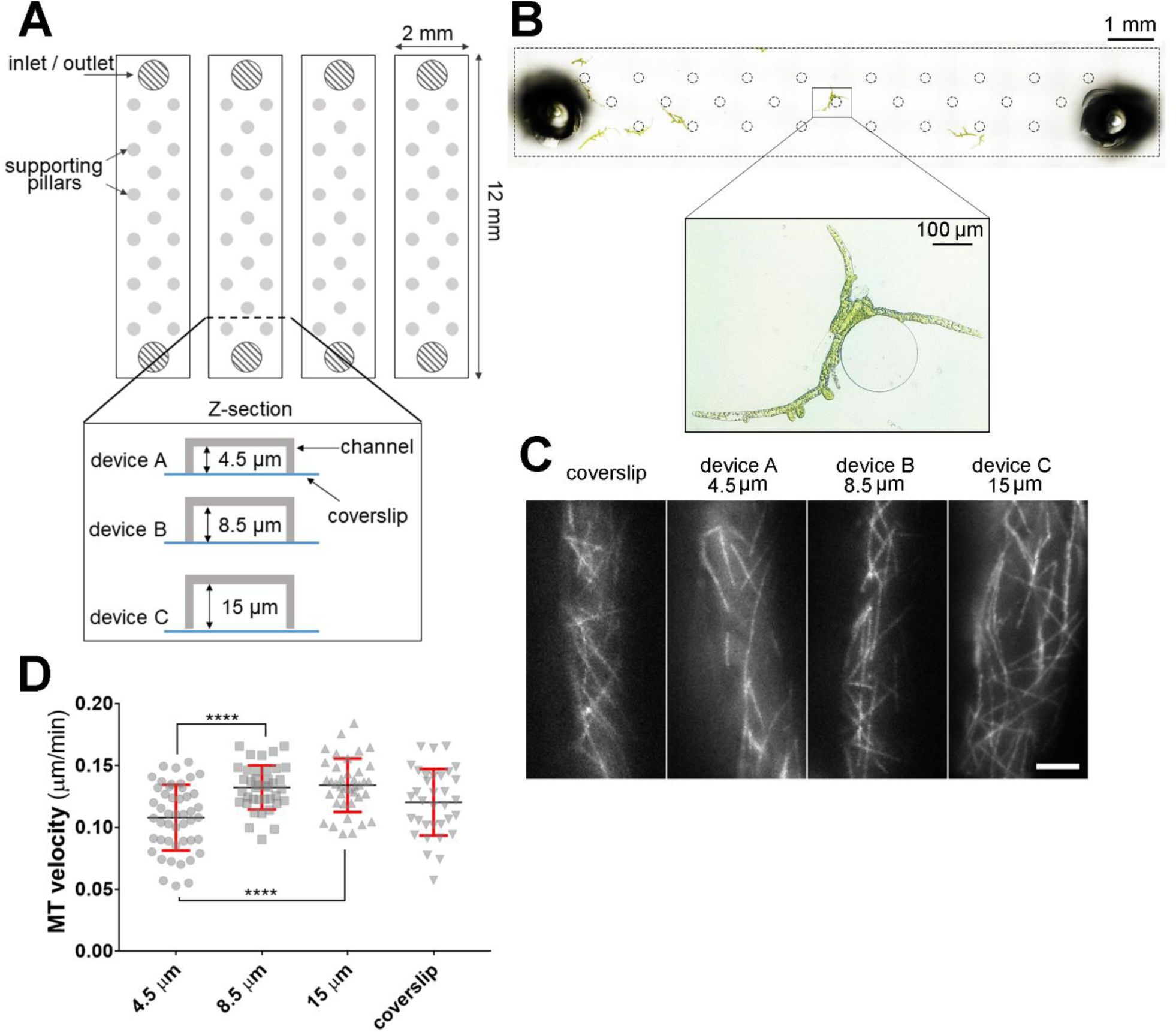
Optimising microfluidic device for HILO live-cell imaging. **(A)** Schematic drawing of the microdevice used for protonema culture and HILO imaging. Three channel depth: 4.5, 8.5, and 15 µm were tested. **(B)** Bright-field image of the single channel (15 µm device) with protonema cells cultured for 3 days. Channel borders and supporting pillars are indicated with dotted black lines. **(C)** Representative images of microtubules HILO imaging obtained in four different setups: coverslip sample, 4.5 µm device, 8.5 µm device or 15 µm device. Scale bar = 10 µm. **(D)** Microtubule velocity (growth rate) slightly decreased in the 4.5 µm channels (*n* = 10, 9, 8 and 6 cells for 4.5, 8.5, 15 µm microdevice and coverslip samples, respectively; mean ± SD; **** *p* < 0.0001 one-way ANOVA).

### 2. Introducing/washing out compounds during live-cell imaging

Microfluidic devices have been successfully used for introducing compounds and subsequent washout in bacteria and animal cells^18–20^; however, unlike individual bacteria or animal cells, the plant cells are connected by cell walls and form irregular shaped clusters (e.g. Figure 1B). Presumably, the plant cells injected into the microdevice can partially block the liquid flow in the channel, which in turn would affect the washout efficiency. Although it is possible to digest the cell wall and isolate single cells (protoplasts), the cell wall plays an important role in plant cell physiology ^21^. Therefore, we aimed to perform the washout experiments without isolating protoplasts.

The device setup for introducing compounds/washout experiments is presented in Figure 2A (sectional diagram) and 2B (a macroscopic photo). In brief, one inlet hole of a microfluidic channel was connected to a syringe pump and the second inlet hole was covered with a drop of growth medium with or without the compound. Initially, we tested the washout efficiency in the channels with growing protonema cells by measuring the fluorescent intensity during introducing/washing the fluorescent dye AlexaFluor 488 at a final concentration 0.5 µM (Figure 2C, D; Supplemental Movie 2). We also measured the background fluorescent levels (Figure 2D ‘background’) prior to introducing AlexaFluor 488 as the reference for washing efficiency. In each experiment, we observed that the fluorescent intensity reached the background level within 90 s (*n* = 8) after start of perfusion (Figure 2D). These results indicate that our washout protocol is robust and efficient even with the protonema cells growing in the device.

**Figure 2.**
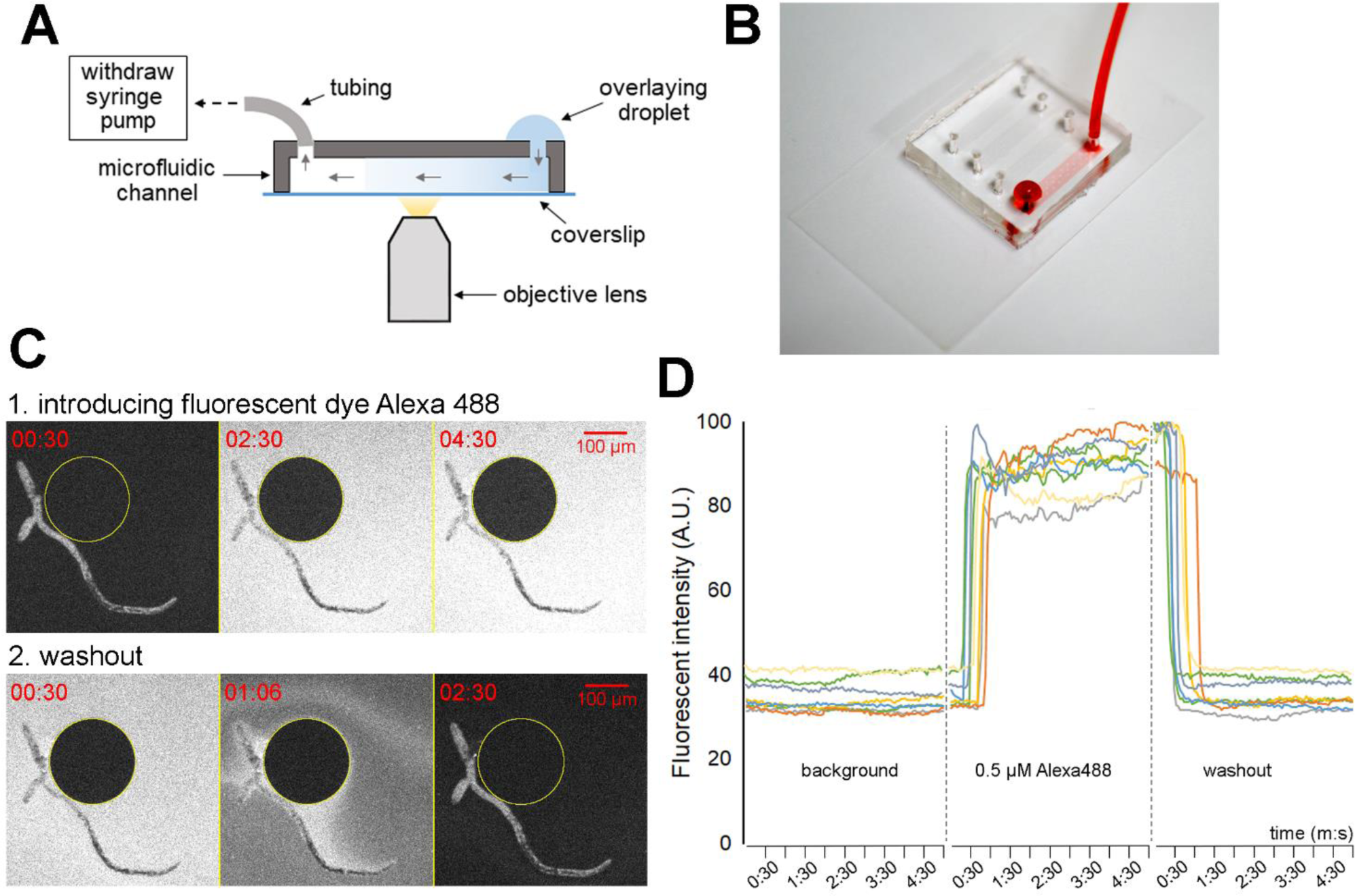
Testing washout setting and efficiency with protonema cells. **(A)** Sectional diagram of the washout setup (not to scale). Outlet hole of the microfluidic channel was connected to the syringe pump with tubing. Inlet hole was covered with a drop of liquid with/without the compound. **(B)** A macroscopic photo of washout setup. Channel is filled with red dye for illustration purposes. **(C)** Representative images of introducing/washing out fluorescent dye AlexaFluor 488 at final concentration 0.5 µM. See also Supplemental Movie 2. **(D)** Changes in average fluorescent intensity (A.U.) in the microfluidic channels before introducing the dye (background), during dye perfusion (0.5 µM AlexaFluor 488) and washout. Fluorescent intensity was measured in five randomly selected areas close to the protonema cells at each time point; mean intensity is plotted on the graph. Each line represents individual experiment (*n* = 8).

### 3. Microtubule nucleation assay

To test if the liquid flow in the channel moves the cells and affects the imaging quality, we conducted a microtubule nucleation assay^22^, using a high-magnification TIRF lens. We used oryzalin treatment at a final concentration of 20 µM for ≥5 min to depolymerise microtubules (Supplemental Figure 1A; Supplemental Movie 3). Next, we washed out oryzalin with the culture medium and observed microtubule re-growth ≥120 s (*n* = 6) after start of perfusion (Supplemental Figure 1A, B; Supplemental Movie 3). The acquired images were of high quality and the observed cells did not drift during imaging. Furthermore, the microtubule density and timing of microtubule nucleation after drug removal (Supplemental Figure 1B) were comparable with the previously published results in the protonema cells^22^. Thus, our microfluidic device can be used for washout experiments combined with high-resolution imaging.

### 4. Observing cytoskeleton behaviour in development

Next, we tested if the present version of the device is suitable for long-term HILO imaging of developmental changes in microtubule arrangement. Some gametophore cells undergo reprogramming and become protonema cells after excision^15^. We speculated that this process would be accompanied by changes in cytoskeletal dynamics, as the microtubule organisation drastically differs between these two cell types. In gametophore cells, microtubules form two-dimensional cortical arrays^23^ typical for majority of vascular plant cells^24^; however, in protonema cells, they are distributed in the cytoplasm in a 3D manner, forming the so-called endoplasmic microtubule network^22,25^. We performed long-term (48 h) HILO imaging to monitor changes in microtubule organisation in the excised gametophore cells injected into the microfluidic device (Figure 3A, B; Supplemental Movie 4). We observed cortical microtubule arrays gradually disorganising, and tip growth and phragmoplast formation in some cells (Figure 3C; Supplemental Movie 5). Furthermore, we estimated changes in the microtubule network by calculating the average microtubule orientation and anisotropy over time (Figure 3D, E) using FibrilTool plugin^26^. Average microtubule orientation changed from being perpendicular to longer cell axis (64° ± 22.8 at time 0, *n* = 8) to more parallel (30° ± 26.8 after 12 h). Anisotropy score (Figure 3E) ranging from 0 (purely isotropic arrays) to 1 (purely anisotropic arrays) reflects the alignment of microtubules. For instance, anisotropy values for parallel microtubule arrays, such as cortical microtubules, typically range between 0.2–0.3^26–28^. In our experiments, the initial values for anisotropy were lower than expected (0.09 ± 0.04 at time 0) and recovered in 3 h (0.2 ± 0.11). Later, anisotropy scores gradually decreased, until they reached similar values for protonema cells (0.12 ± 0.04 after 12 h vs. 0.17 ± 0.06, *n* = 8 for excised gametophore cells and *n* = 7 for protonema cells, respectively). Interestingly, we observed that microtubule organisation changed in all gametophore cells, regardless of whether we observed regeneration in protonema cells.

**Figure 3.**
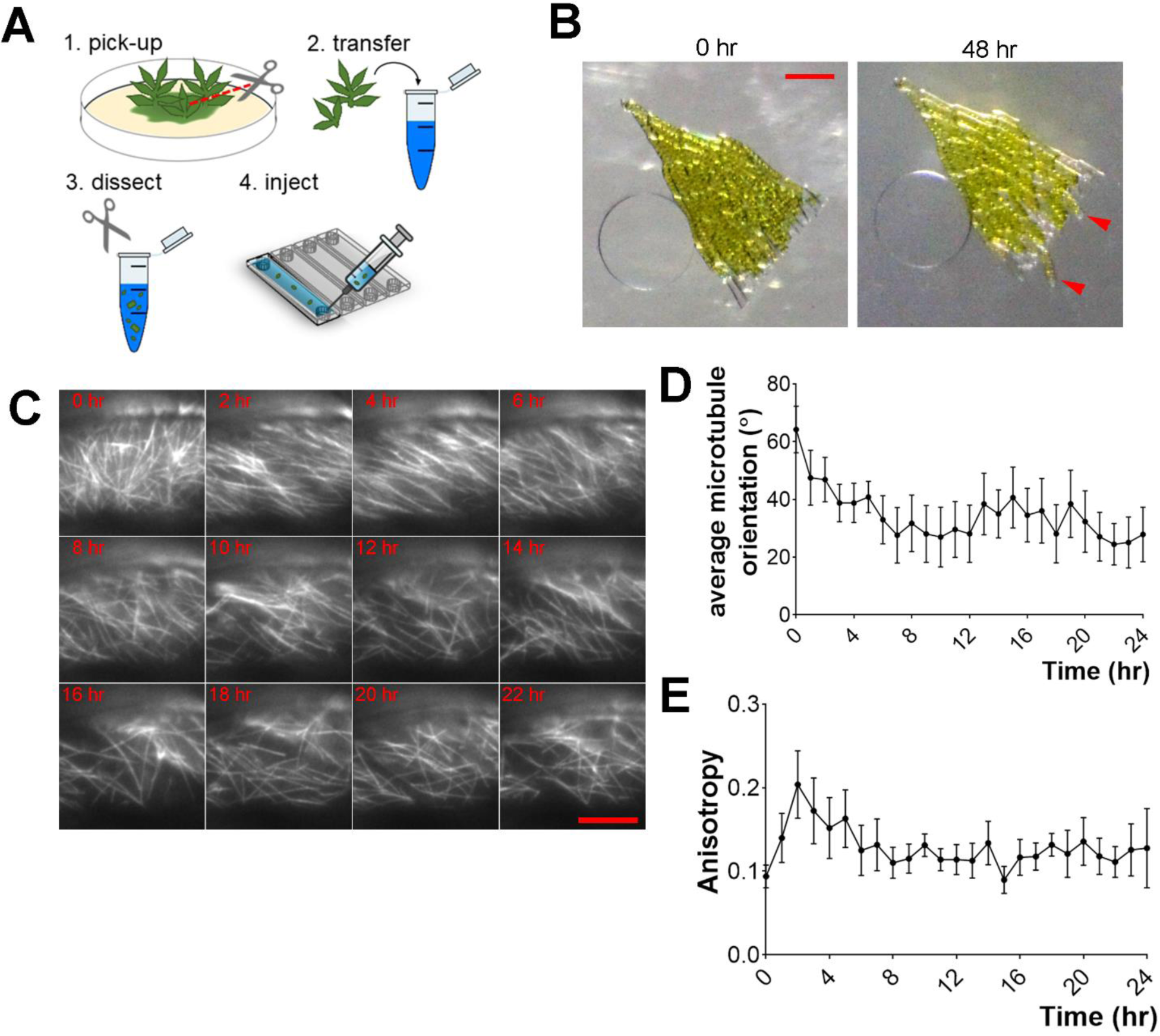
Cytoskeleton remodelling during gametophore regeneration observed with long-term HILO imaging. **(A)** Flowchart for gametophore excision and injection in the microfluidic device. **(B)** Representative image of excised gametophore cells injected in the microdevice and protonema regeneration (red arrowheads) after 48 h. Sample was maintained in 1-min dark/4-min light conditions for imaging. Scale = 100 µm. See also Supplemental Movie 4. **(C)** Representative images of microtubule dynamics in excised gametophore cells. Gradual disassembly of cortical microtubule arrays can be seen (e.g. 4 and 12 h). Scale = 10 µm. See also Supplemental Movie 5. **(D)** Graph of average microtubule orientation (angle) during 24 h time-lapse (*n* = 8, mean ± SEM). Measured every 1 h with FibrilTool ImageJ plugin. **(E)** Graph of anisotropy (score between 0 and 1) changes during 24 h time-lapse (*n* = 8, mean ± SEM). Measured every 1 h with FibrilTool ImageJ plugin.

## Discussion

Over the past few years, microfluidic technology is slowly gaining popularity among plant researchers^2^. Microfluidics has been successfully applied to study root growth^10^ and interaction with bacteria^24^, pollen tube attraction^30^, embryo live-cell imaging^31^ and protoplast fusion^32^. Although we have already applied microfluidic technology to perform HILO imaging^33^, here we conducted additional tests to determine the optimal conditions and present several new applications of the microfluidic device, such as washout and long-term imaging. By trying several versions of the microdevice, we found that 15 µm deep channels are sufficient to increase the visible cell surface for HILO imaging. Furthermore, we analysed the microtubule growth rate (velocity) as a representative parameter of microtubule dynamics^34^ and demonstrated that mild space restrictions did not cause significant changes in the microtubule growth rate. We suggest that 15 µm deep microdevice can be applied to live-cell imaging of moss *P. patens* in conditions close to natural.

One of the major applications of microfluidics in cell biology is the precise spatiotemporal control of liquid in the channels. It can be used to supply fresh growth medium, create chemical gradients, or supply/remove compounds. The latter application was tested in the present study. We demonstrated that washout is robust and efficient with few clusters of protonema cells growing in the channel. Importantly, we never observed cells drifting during liquid perfusion; therefore, high-resolution imaging could be performed while introducing/washing compounds (e.g. microtubule nucleation assay, Supplemental Figure 1). Since plant cells can be cultured in the microdevice for several days, long-term drug treatment combined with imaging is also feasible. Although, here we tested only HILO imaging, we suggest that our microdevice can be also used for washout assays during high-resolution confocal microscopy, for example, to image cell division or organelle movements.

Furthermore, we confirmed that microfluidic device enables long-term HILO imaging of plant samples. As a proof-of-concept, we imaged microtubule reorganisation associated with regeneration of excised gametophore cells^15^ in the microdevice. It is possible that the gametophore excision or injection in the microdevice transiently disrupts the microtubule arrays, based on low anisotropy values (Figure 3E); however, the anisotropy score was eventually recovered to expected values for cortical microtubules^26–28^. Previous study demonstrated that gametophore cells regenerating as protonema produce certain signals, inhibiting protonema regeneration of the surrounding cells^15^. Interestingly, we observed changes in the microtubule organisation, such as disassembly of cortical arrays, in all gametophore cells. This suggests that all gametophore cells undergo reprogramming and dedifferentiation; however, tip growth and cell division are inhibited in a subset of the cells.

Traditionally, to perform HILO or TIRF imaging, a part of the plant tissue (e.g. root, leaf or protonema cells) is excised and flattened between two coverslips^6,22^. Some cells may be damaged during sample preparation, which in turn can affect the obtained results. Furthermore, due to poor gas exchange through the coverslip, a time-frame available for imaging is limited. For instance, in our experience, the protonema cells placed between coverslips are viable for approximately 30 min. Therefore, we consider using the microfluidic device for HILO imaging as a considerable improvement over the existing methodology. Initially, the cells can be cultured for several days in the microdevice after injection, permitting adaptation to the new environment. Microdevice is fabricated from a gas-permeable polymer that has been proven as biocompatible for both plant^9^ and animal cells^11,12^. According to our data, shallow microfluidic channels are as efficient as coverslips to flatten cells for HILO imaging. Furthermore, long-term HILO imaging can be performed using the microdevice. With minor modifications, such as changing channel depth, the device developed in this study would also be suitable for HILO imaging in other model plant systems (e.g. *Nicotiana tabacum* BY-2 and *Arabidopsis thaliana* culture cells). We speculate that application of the microfluidic device to HILO microscopy and inhibitor assays will provide valuable insights into plant cell and developmental biology.

## Methods

### Moss culture and transformation

We generally followed the protocols described by Yamada et al^35^. *Physcomitrella patens* culture was maintained on BCDAT (abbreviation stands for stock solutions <u>B</u>, <u>C</u>, <u>D</u> and <u>A</u>mmonium <u>T</u>artrate) medium at 25 °C under continuous light. Transformation was performed with the polyethylene glycol-mediated method, and successful cassette integration was confirmed by PCR^35^ and microscopy. We used the GH line^36^, expressing <u>G</u>FP-tubulin and <u>H</u>istoneH2B-mRFP for microtubule HILO imaging and washout experiments. To observe cortical microtubules in gametophore, we created a new line, namely GGH, comprising the <u>G</u>CP4promoter-<u>G</u>FP-tubulin and <u>H</u>istoneH2B-mRFP. *P. patens* lines, used in this study are described in Supplemental Table 1.

### Plasmid construction

Vectors and primers used in this study are described in Supplemental Table 2. GCP4 promoter-GFP-tubulin vector, assembled with In-Fusion enzyme (TAKARA Clontech), contains eGFP-PpTuA^37^ under endogenous GCP4 promoter (2 kb upstream of GCP4 gene; accession number Pp3c14_20420, Phytozome) and G418 resistance cassette for selection and homologous recombination sites targeting homeobox 7 (hb7) gene locus^38^. For <u>H</u>istoneH2B-mRFP we used previously created vector pGG616^36^, comprising human histone H2B-mRFP fusion gene, directed by the E7113 promoter^39^.

### Microdevice fabrication

The mold for poly-dimethylsiloxane (PDMS) device was prepared by spin-coating negative photoresist (SU-8 3005 for 4.5 and 8.5 µm deep device, 3010 for 15 µm deep device; Microchem Corp.) on a silicon wafer. Maskless lithography system (DL-1000; Nano System Solutions, Inc.) was used to create microchannel designs in the photoresist layer. To make the PDMS devices, a pre-polymer mixture of PDMS (Sylgard 184; Dow Corning) was prepared by mixing the elastomer base and curing agent at a ratio of 10:1 and pouring onto the mold. After degassing for 30min, the mold was placed in an oven (65°C) for 80min. The inlet to the microchannel was created using a biopsy punch (1.5 mm diameter; Harris Uni-Core). The PDMS device and a coverslip (Matsunami, glass thickness 0.16–0.19 mm) or glass bottom dish (Matsunami; dish diameter 35 mm; glass diameter 27 mm; glass thickness 0.16–0.19 mm) were both exposed to air plasma for 50 s, pressed together, and heated for 2 h at 70 °C for irreversible bonding of PDMS layer on the glass surface.

### Introducing moss in the microdevice

Microdevice inlet holes were connected to the small pieces of polyethylene tubing (I.D. 0.76 mm; O.D. 1.22 mm; Intramedic). Microdevice was exposed to vacuum for 15–20 min and immediately filled with BCDAT liquid medium^35^ using a 1 ml syringe with needle (Terumo, 0.80 × 38 mm). This step helps to prevent air bubbles in the channels. Next, the microdevice was sterilised under the UV (2000 × 100 µJ/cm^2^; UVP Crosslinker CL-1000, AnalytikJena). Protonema cells of *P. patens* were grown on cellophane for 5–7 days^35^. We collected the cells from approximately 1/16 of standard Petri dish and sonicated in 5 ml of BCDAT liquid medium^35^ (without agar). The sonicated cells were filtered through 50 μm nylon mesh to isolate small cell clusters (typically 1–4 cells), and the flow-through fraction was gently injected in the microdevice. Microdevice with protonema cells was submerged in the BCDAT medium to prevent liquid evaporation from the channels; the dish was sealed with parafilm and left for 3–4 days at 25 °C under continuous light. For gametophore imaging, we picked several gametophores from the moss colonies cultured on the BCDAT plates, dissected them into small pieces using scissors in the BCDAT liquid medium and injected in the microdevice using 1 ml syringe (see Figure 3A). Imaging was conducted immediately after sample preparation. We recommend using young gametophores, containing 3–4 leaves, for this experiment.

### Introducing compounds/washout experiments

For washout experiments, one access hole (outlet) was connected to the syringe pump (YMC, YSP-202) using polyethylene tubing (I.D. 0.76 mm; O.D. 1.22 mm; Intramedic) and the second access hole (inlet) was covered with a small drop of BCDAT medium with or without drug (see Figure 2A, B). As the BCDAT medium may contain inorganic salt precipitates, we passed it through 20 µm filter just prior to the experiments. The remaining holes were covered with small pieces of scotch tape to prevent liquid cross-contamination. The device was then mounted on the microscope stage and additionally fixed with a scotch tape. Introducing compounds/washouts were performed in the ‘withdraw’ pump mode at a stable flow rate of 10 µl/min. Every 2–3 min, liquid was supplied on top of the inlet hole using a 20 µl pipette. Notably, connecting tubing may damage the cells located near the outlet hole; therefore, we suggest to image cells growing in the middle of the channel.

### Live-imaging microscopy and data analysis

Preparation of the coverslip samples was performed as previously described^22^. Epifluorescence microscopy to test the washout efficiency was conducted with Nikon’s Ti microscope (10× 0.45-NA lens), equipped with an electron-multiplying charge-coupled device camera (Evolve; Roper). HILO imaging was performed with a Nikon Ti microscope with a TIRF unit, a 100× 1.49 NA lens and EMCCD camera (Evolve; Roper). All imaging was performed at 24–25 °C in the dark, except for the gametophore regeneration assay that requires light (4-min light/1-min dark cycle). The microscopes were controlled by the Micro-Manager software and image data were analysed with ImageJ. Microtubule density in the microtubule nucleation assay was evaluated manually. We used FibrilTool ImageJ plugin^26^ to calculate the average orientation of microtubules and anisotropy in the given region of interest (ROI) from raw images.

## Acknowledgement

We would like to thank Dr. Naoki Yanagisawa for initial device design, advice on washout experiments, comments and discussion on the manuscript; Shu Yao Leong for providing GGH line, Dr. Moe Yamada, Dr. Peishan Yi and Shu Yao Leong for comments on the manuscript. This work was funded by JSPS KAKENHI (17H06471) to G.G.

---

**Supplemental Figure 1.**
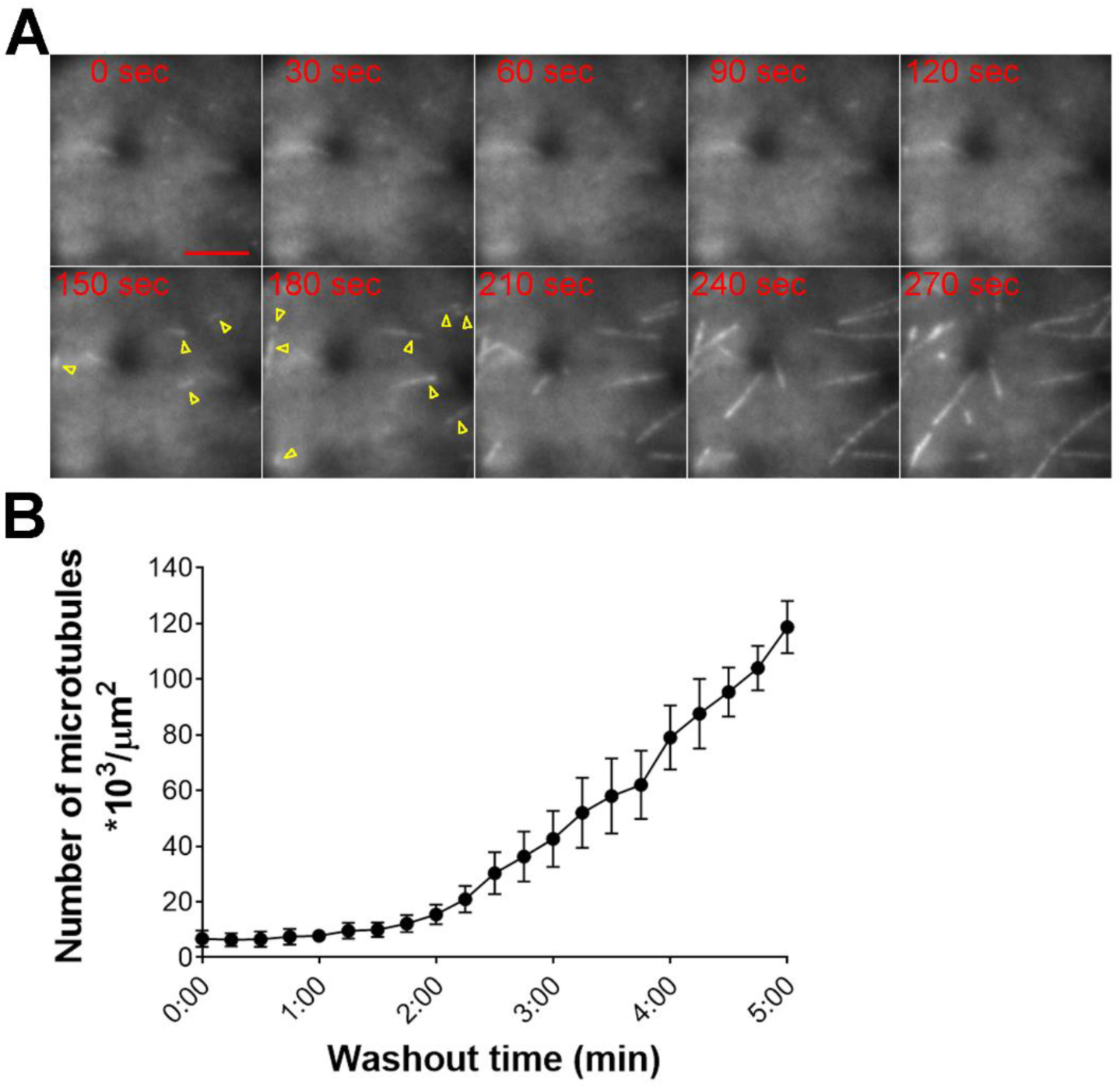
Microtubule nucleation assay in the microdevice. **(A)** Microtubule nucleation observed after oryzalin washout. New microtubules are presented with yellow arrowheads. Oryzalin washout commenced at time 0. Scale = 5 µm. See also Supplemental Movie 3. **(B)** Microtubule density (number of microtubules per area µm^2^) was manually evaluated every 15 s from the washout commencement (*n* = 6, mean ± SEM).

**Supplemental Table 1.** Transgenic moss lines used in this study.

| Line# | Clone# | Name | Plasmids | Resistance | Reference |
| --- | --- | --- | --- | --- | --- |
| 0002 | 3 | **GH** (GFP-tubulin/H2B-RFP) | GFP-tubulin<br>pGG616 | Zeocin, G418 | Nakaoka et al., 2012 |
| 0704 | 15, 30 | **GGH** (GCP4promoter-GFP-tubulin/H2B-RFP) | pSY100<br>pGG616 | Zeocin, G418 |  |

**Supplemental Table 2.**
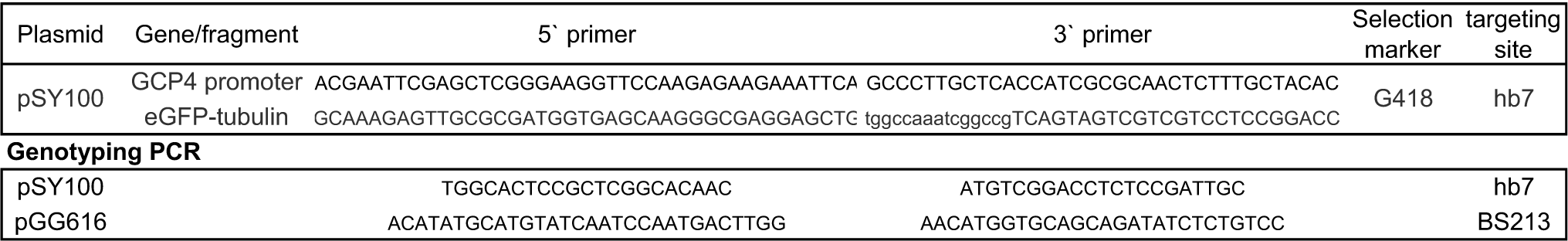
Primers and plasmids used in this study.

**Supplemental Movie 1.** HILO imaging of microtubule network using different set-ups: coverslip sample, 4.5, 8.5 or 15 µm deep microfluidic channels. Imaging was conducted in the protonema cells expressing GFP-tubulin. Images depict a single focal plane acquired every 3 s.

**Supplemental Movie 2.** Introducing fluorescent dye AlexaFluor488 *(left)* and subsequent washout *(right)*. Position of the supporting pillar is indicated with *yellow* circle. Protonema cells are expressing GFP-tubulin. Images depict a single focal plane acquired every 3 s.

**Supplemental Movie 3**. Microtubule nucleation assay using microdevice. *(left)* introducing microtubule-depolymerising drug oryzalin at final concentration 20 µM, *(right)* oryzalin washout and microtubule regrowth. Imaging was conducted in protonema cells expressing GFP-tubulin. Images depict a single focal plane acquired every 3 s.

**Supplemental Movie 4.** Regeneration of excised gametophore cells to protonema in the microdevice. Imaging was conducted in protonema cells expressing GFP-tubulin under native GCP4 promoter (microtubules, green) and HistoneH2B-mRFP (nucleus, magenta). Images depict a single focal plane acquired every 5 min.

**Supplemental Movie 5.** HILO imaging of microtubule network in excised gametophore cells during regeneration. Imaging was conducted in protonema cells expressing GFP-tubulin under native GCP4 promoter. Phragmoplast can be seen at 25 h 40 min time-frame. Images depict a single focal plane acquired every 5 min.

